# Personalized genealogical history inferred from biobank-scale IBD segments

**DOI:** 10.1101/2019.12.19.883108

**Authors:** Ardalan Naseri, Kecong Tang, Xin Geng, Junjie Shi, Jing Zhang, Xiaoming Liu, Shaojie Zhang, Degui Zhi

## Abstract

When modern biobanks collect genotype information for a significant fraction of a population, dense genetic connections of a person can be traced using identity by descent (IBD) segments. These connections offer opportunities to characterize individuals in the context of the underlying populations. Here, we conducted an individual-centric analysis of IBDs among the UK Biobank participants that represent 0.7% of the UK population. On average, one UK individual shares IBDs over 5 cM with 14,000 UK Biobank participants, which we refer to as “cousins”. Using these segments, approximately 80% of a person’s genome can be reconstructed. Also, using changes of cousin counts sharing IBDs at different lengths, we identified a group, potentially British Jews, who has a distinct pattern of familial expansion history. Finally, using the enrichment of cousins in one’s neighborhood, we identified regional variations of personal preference favoring living closer to one’s extended families. In summary, our analysis revealed genetic makeup, personal genealogical history, and social behaviors at population scale, opening possibilities for further studies of individual’s genetic connections in biobank data.

## Introduction

An individual’s genealogical history is often of interest to the innate curiosity about one’s ancestry. However, it has been challenging to collect complete and accurate family pedigree of an individual. Further, the pedigree record is often incomplete and lacking objective, quantitative details of one’s ancestors and relatives. As a result, population studies of personal genealogical history are often void of quantitative rigor.

Thanks to the establishment of modern biobanks and direct-to-consumer (DTC) genetic testing companies which contain genotype information of a relatively large fraction (0.1%-5%) of a population, it is now possible to identify dense genetic connections among individuals. This opens the possibility to study the quantitative details of one’s genealogical history and identify emerging patterns.

Here, we present the first comprehensive analysis of the personal genealogical history inferred from all identity-by-descent (IBD) segments of an individual in a biobank of a large modern population. We first made a high quality call set of IBD segments over 5 cM among all 500,000 UK Biobank ^1^ participants. Based upon the genetic connections revealed through these IBD segments, we investigate the “cousin cohort”, all persons who share at least one IBD segment, of individuals. We discuss a set of genealogical descriptors (GPs) based on the cousin cohort and show that these genealogical descriptors offer very rich information about one’s genetic makeup, personal genealogical history and social behaviors.

## Results

### IBD segment calling and quality assessment

Traditional methods for IBD segment identification are not scalable to data sets containing millions of samples. Using RaPID ^2^, an efficient and accurate method, we identified 3.5 billion IBD segments over 5 cM within the 22 autosomes among 487,409 UK Biobank participants (**Methods: IBD segment calling using RaPID**). Thanks to the efficiency of RaPID, we achieved this in 5.25 days using a single core CPU with 6.34G peak memory. This translates to, on average, each person having 5 cM genetic connections to 14,000, or about 3% of UKBB persons, whom we call genetic cousins or cousins for short. These segments offer on average 10x coverage of the overall diploid genome for a UK individual.

Noting that this density of IBD segments is higher than that reported by existing studies ^3^, we took extra caution assessing the quality of our results. Since quality assessment of IBD segment calls of a large cohort is a less-studied problem, we developed the following strategies: First, we compared the kinship coefficients derived from RaPID’s IBD calls against a standard genotype-based relatedness caller, KING ^4^. We verified that RaPID’s call is indeed consistent with the theoretical expectation for close relatives (Supplementary Figure S1). Notably, IBD segments provide much deeper information about realized genetic relationships than KING’s results. For example, only 50% of the IBD segments of length 70-80 cM identified by RaPID belong to the public release ^1^ of 3rd degree or closer relatives identified by KING (Supplementary Figure S2).

Second, we verified the consistency of RaPID’s calls with traditional methods. As not all methods can run to call >5 cM IBD segments for all UK Biobank participants without extravagant computing resources, we used a subset of 200,000 participants and ran over the chromosome 22. We ran traditional methods including GERMLINE ^5^, Refined IBD ^6^, and also a recent method iLASH^7^, and calculated the overlap between these sets of IBD calls. We found that RaPID’s results included almost all segments called by other methods: 95.8% of GERMLINE’s calls, 97.2% of iLash’s calls, and 99.4% of Refined IBD’s calls (Supplementary Table S1). Interestingly, RaPID detected 16.3% more segments that were missed by all other methods.

Third, we leveraged the monozygotic (MZ) twin pairs to estimate the detection power of IBD segments. While MZ twins should match their entire chromosomes, in real data, the long IBD segments are interrupted by switch errors in phasing. Fortunately, these MZ twins are easy to call from non-IBD calling methods, such as KING. Using the 179 twin pairs identified by KING, we observe that using the average haplotype matching identity over the moving average of windows with 100 SNPs, the switch errors are notably visible (Supplementary Figure S4). Thus, these switch errors created IBD segments of various lengths. A perfect IBD segment detector should identify multiple segments that in aggregate cover the entire length of the genome.

We define the power of a method over one sample as the percent of the sites over the entire genome that was covered by any segment detected by the method. The average power values over all twin pairs of these methods are shown in Supplementary Table S2 and Supplementary Figure S5. RaPID has consistently demonstrated higher power when compared to other methods. It is also of interest that the power values of all methods between British twin pairs are higher, probably due, in part, to the fact that British individuals have superior genotyping and phasing quality. Overall, RaPID’s power is approximately 1% higher than GERMLINE for British and 5% higher for non-British.

Fourth, we leveraged parent/offspring pairs to evaluate the accuracy of IBD calling. Overall, each method achieved accuracy over 99.9%, as there is very little chance that a false positive segment over 5 cM is called by any of these methods. The quality of IBD calls made by RaPID is high according to the aforementioned quality assessment. We thus proceeded to the downstream analysis of the IBD calls.

### Cousin count distribution by UK regions (county)

Overall, average kinship coefficients across self-reported ethnicities are consistent with expectations (Supplementary Figure S6). The distribution of cousin counts of all UKBB participants is shown in Figure 1a. While the peak with cousin count < 4000 is mainly due to minorities, there are two peaks noticeable for the British people (Supplementary Figure S7).

**Figure 1:**
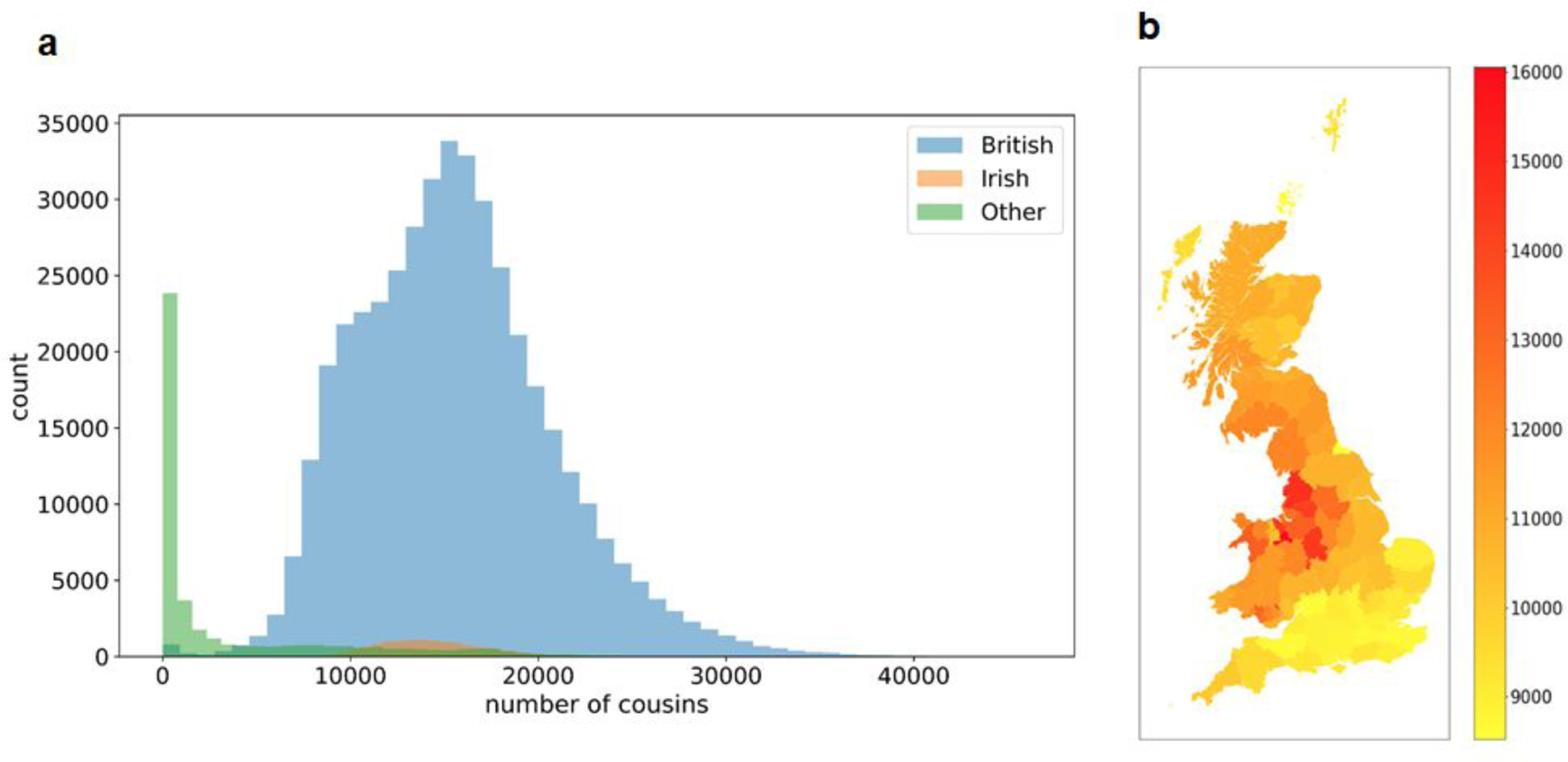
Cousin count distribution (unadjusted stacked histograms of British, Irish, and others). (a) Overall distribution over all individuals of all ethnicities. (b) Average cousin counts of all UK areas (except Northern Ireland as home location of Northern Ireland is not available).

We found that individuals from central England (e.g., Manchester) have almost twice as many cousins as those from southern England (e.g., London) (Figure 1). The two peaks persist even after adjusting the cousin counts by the uneven sampling rates of different regions (**Methods: cousin count adjustment by regional sampling rate**). Also, the inter-region cousin counts indicating clusters that are consistent with geographic boundaries (Supplementary Figure S8). It is evident that Welsh and Scottish people form relatively tight clusters, respectively. Northern England has a visible cluster but also has significant sharing with Scottish individuals. Southern England seems to be less a cluster and share IBDs evenly to most other regions.

### Personalized genealogical descriptors

Using all cousins of an individual as a cohort, we can infer the personalized genealogical history of each individual. This is only possible as each participant has a sufficiently large number of cousins (estimated 14,000) on average. Besides the obvious choice of total cousin counts, we developed the following three sets of genealogy descriptors (GPs): the genome-wide coverage of IBD segments, the cousin counts stratified by IBD segment length, and the cousin enrichment within a neighborhood.

#### 5 cM IBD segments cover the entire diploid genome of an average UK individual 10 times

While 5 cM IBDs on average cover each individual’s genome 10 times, the coverage is highly uneven. To capture the individual variability of genome coverage, we used 5 genealogical descriptors: percent genome covered by IBD segments from 1, 2-5, 6-10, 11-20, 21-50, or larger than 50 cousins. Our analyses are focused on British people as they are well-represented (the results for all ethnicities are available at Supplementary Figure S9). We found that this coverage is highly uneven among British individuals (Figure 2a). Overall, 50% of the British have over 85% of their genome, and 75% of British individuals have over 80%, covered by at least 1 IBD segment. For coverage of 10 IBD segments, 50% British have over 50% covered by at least 10 segments. To an extreme, 5% of the British population (20,000 in UK Biobank) have 20% of their genome covered by over 50 segments.

**Figure 2:**
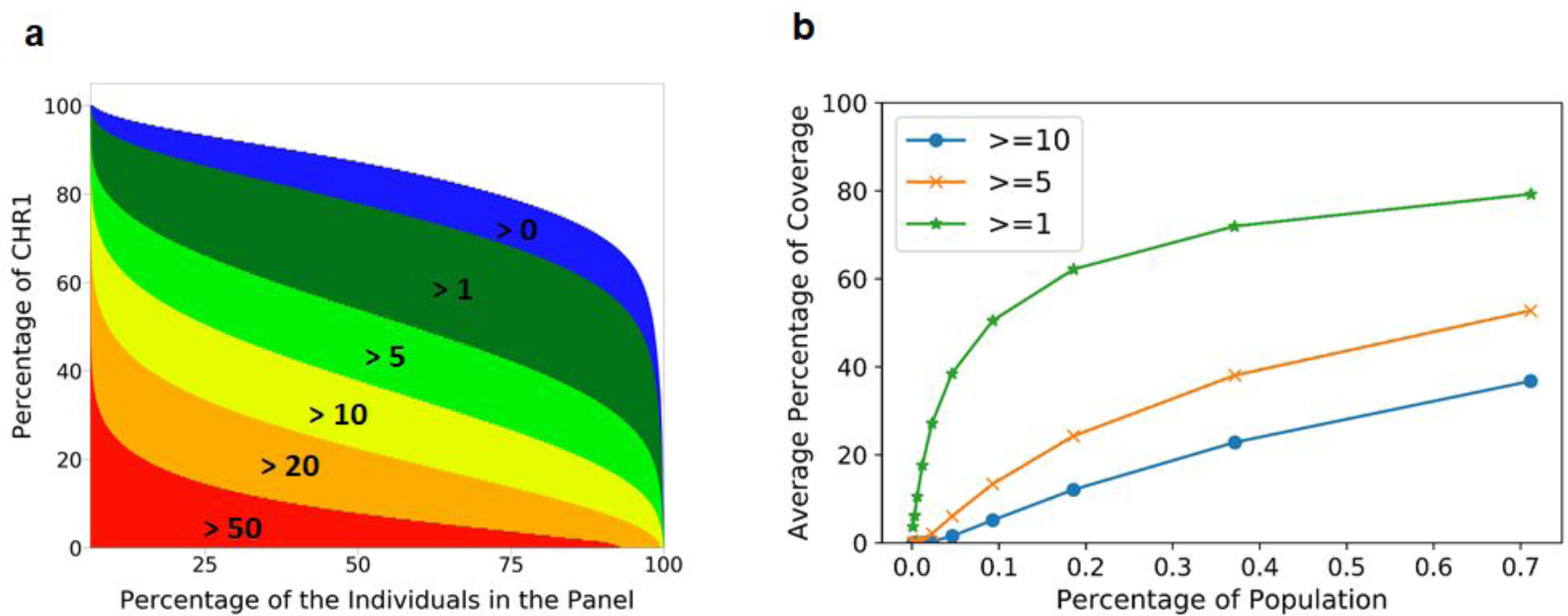
Percentage of genome covered by >5 cM IBD segments in British people in the UK Biobank. (a) Percentage coverage of chromosome 1 by increasing the number of IBD segments, across all British individuals. (b) Average percentage coverage of chromosome 1 by IBD segments.

Further, we investigated the average percent of IBD covered genome (by any IBD segment) as a function of the percent of the population being studied (Figure 2b). At the 0.7% sampling rate of UK Biobank, 80% of the genome is covered by at least 1 IBD segment. Even at 0.1% sampling rate, the average genome coverage remains 50%. This fact implies a much more reliable and base-pair resolution imputation or construction of one’s genome with a very small portion of genome polymorphism (e.g. microarray genotype data) when a biobank scale database is available. Public awareness of this genome reconstructability is needed as to the potential issue of genome privacy (see **Discussions**).

#### Individuals’ cousin count stratified by IBD length reflects familial expansion history

The decay pattern of cousin counts with increasing IBD length reflects population history ^8,9^. Instead of analyzing the gross pattern over all individuals, we analyze the pattern pertaining to each individual. We derived the following cousin counts for each individual: the count of cousins sharing 5-10 cM, c5, and the count of cousins sharing >10 cM segments, c10, also their ratio c=c5/c10. For most of the British individuals, the ratio is centered around 0.13. Notably, there is a small fraction of people, n=1719, with c>1 (Figure 3a). Note that their c5 is among the smallest, and their c10 is among the largest (Figure 3b). Therefore, they represent a population undergoing rapid recent expansion. These individuals are enriched in Greater London, Greater Manchester, and the outskirts of Glasgow (Figure 3c). Also, their population frequency is about 0.4%. All these characteristics match that of British Jewry^10^.

**Figure 3:**
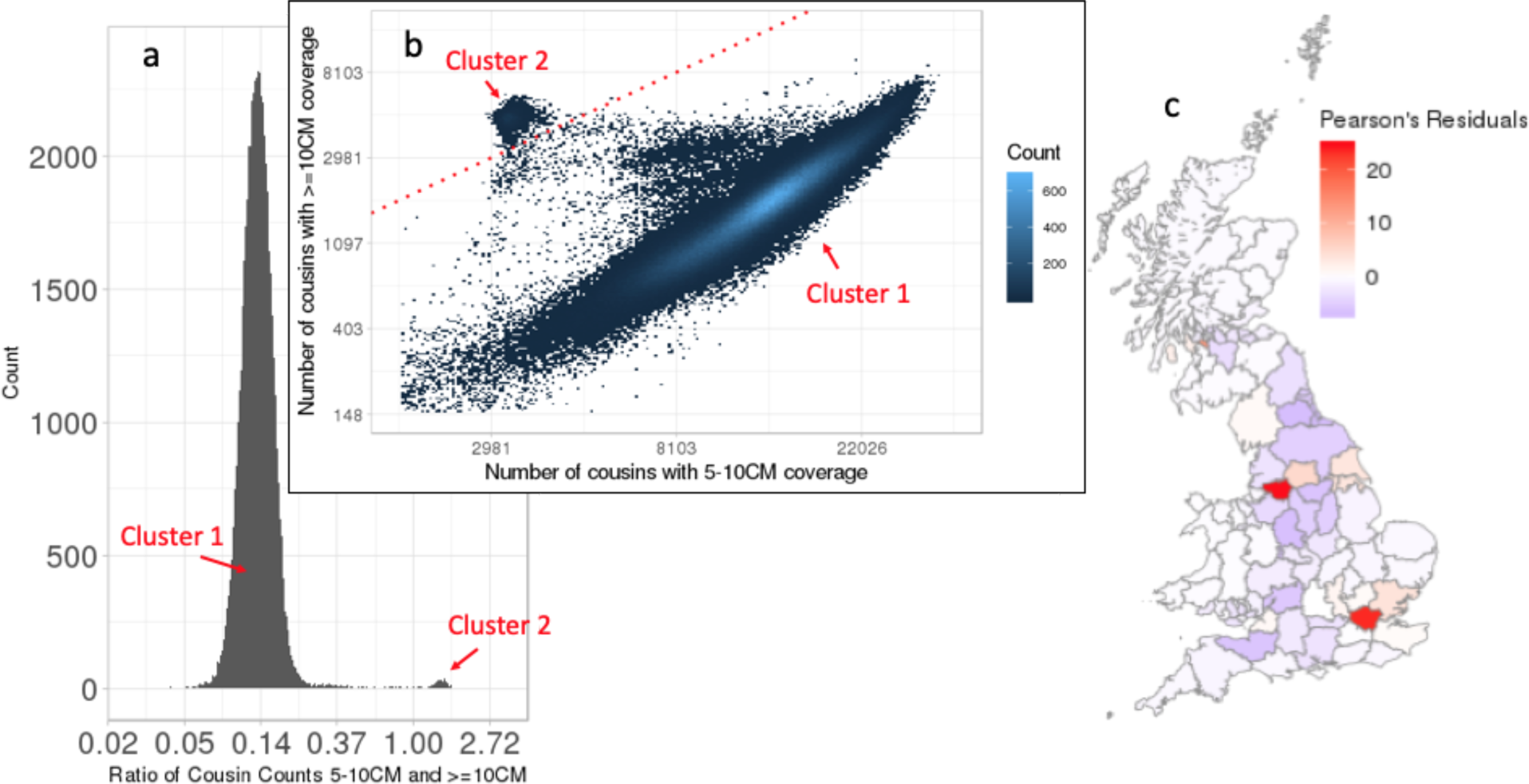
A group of 1719 individuals with distinct IBD patterns. (a) Two clusters are distinctive in histogram of ratio of cousin counts sharing 5-10 cM and sharing 10 cM IBD segments. (b) People in Cluster 2 have low cousin counts in 5-10 cM segments and has high cousin counts in >10 cM segments. Red line is the diagonal. (c) Regional enrichment of Cluster 2.

In order to validate this hypothesis, we downloaded genotype data of an Ashkenazi Jewish individual and phased them using his parents’ genotypes. The trio’s data were downloaded from the Personal Genome Project^11^. We then searched for two phased haplotypes of the son (in Chromosome 1) in the UKBB with a minimum target match length of 200 SNPs, corresponding to about 8 cM, using PBWT-Query ^12^. A detailed description of the pipeline can be found in the Methods section. The first haplotype had 293 hits where 103 of them were in the subset of individuals with high *c* value. The second haplotype had 257, out of which 88 were in the subset of individuals with high *c* values. The fact that over ⅓ of hits were from this 0.4% population suggests that this cluster of high *c* values identifies probable British Jews (p-value<10^−16^, Chi-squared test).

#### Analysis of cousin enrichment in neighborhood reveals regional patterns of preference for living closer to relatives

It is generally expected that an individual lives near to one’s extended family. Assuming cousins represent one’s extended family, we can study the social behavior of individuals. To quantify the preference of an individual towards living closer to their extended family, or local connectivity, we calculated the cousin enrichment within the neighborhood (CEIN). This can be defined as the ratio between the cousin density in the neighborhood to the cousin density across the entire UK. A CEIN of one indicates no preference of local connectivity, and a larger CEIN indicates stronger preference. We confirmed that overall cousin density is indeed enriched by almost 1.7 fold in the neighborhood of a 1km radius of a person. This enrichment showed a sign of slow decay as the neighborhood radius became greater (Supplementary Table S3). CEIN as a measure of local connectivity is quite noisy as indicated by its broad distributions (Supplementary Figure 10). We chose to use cousin enrichment within 25km radius (e25) as the CEIN measure as it is including a greater number of individuals, and thus is numerically more stable.

Interestingly, local connectivity is inversely correlated with 25km neighbor count (Figure 4). Based on e25 we identify 3 clusters: people with weak, moderate, or strong local connectivity. Using the number of neighbors within 25km as a proxy of the living environment, we divided the individuals as dwellers of “Metropolitan”, “Big City”, “Mid-sized City”, “Small City” and “Rural”(**Methods: Designating types of living environment by counting neighbors**). For people in Metropolitan, i.e., neighbor count > 40,000, there is a lack of local connectivity. For people in Big City and Mid-sized City (with the neighbor count in 21,000-40,000), there is a second subgroup of people with e25=1.7. For people with a neighbor count <21,000, the cousin enrichment in the neighborhood is very high, suggesting people living in less-populated areas tend to have higher enrichment of cousins within the neighborhood.

**Figure 4:**
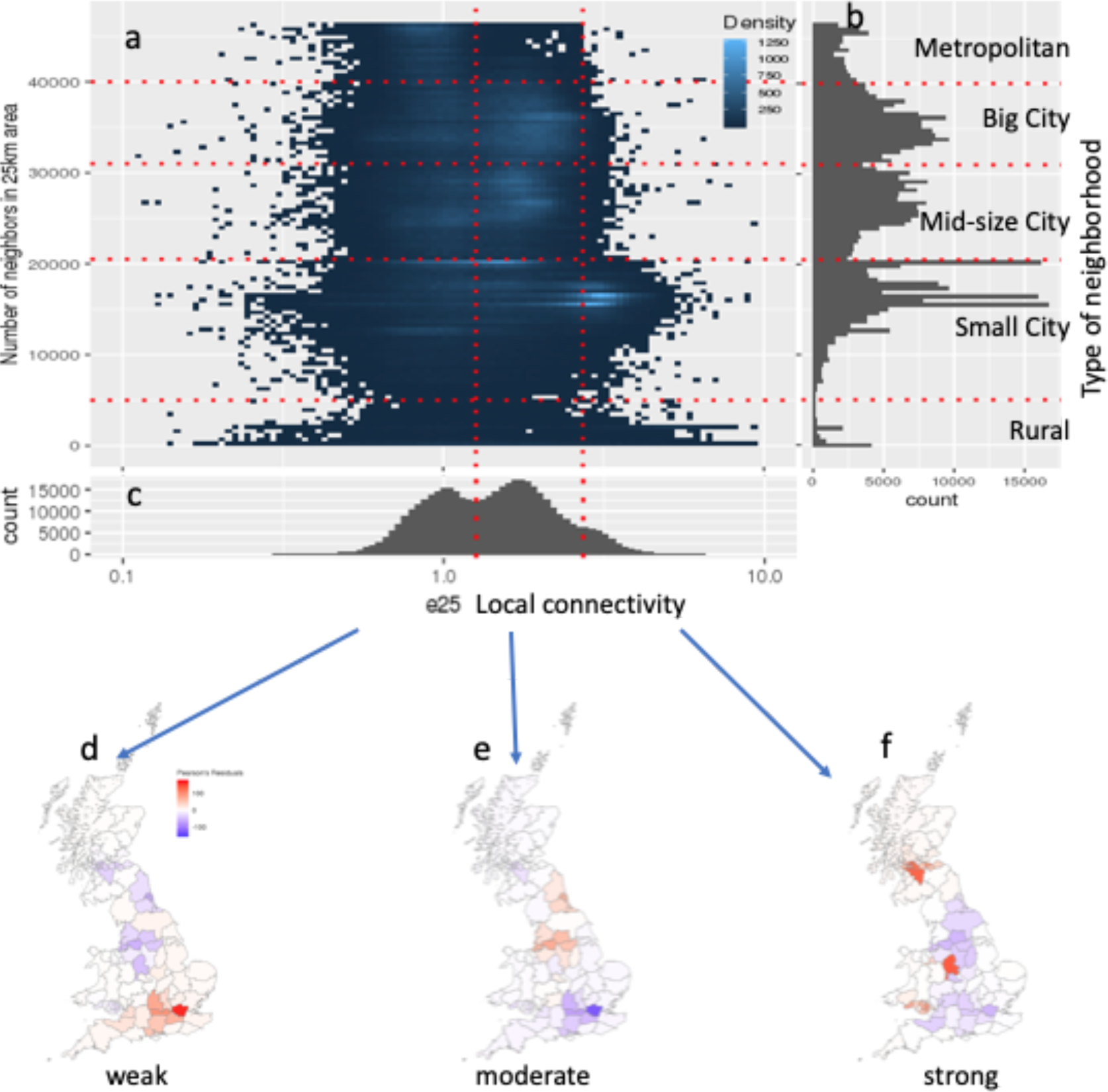
Preference of local connectivity as measured by cousin enrichment in the neighborhood (CEIN). (a) Preference of local connectivity (enrichment of cousins in 25km neighborhood, e25) is overall inversely correlated with population density (neighbor count). (b) Type of neighborhood of an individual can be captured by the number of neighbors in 25km radius. (c) According to enrichment of cousins in 25km radius, UK Biobank participants can be clustered to three groups: people with preferences of weak, moderate or strong local connectivity. (d)(e)(f) maps of regional enrichments of people of weak, moderate, and strong local connectivities.

We further depict the regional differences of people with weak, moderate and strong local connectivities (Figure 4(d)(e)(f) and Supplementary Table S4). Interestingly, the counties that are with highest enrichment for people with weak local connectivities are Greater London, Berkshire, and Oxfordshire, while the counties with highest enrichment for people with strong local connectivity are Staffordshire, Greater Glasgow (South Lanarkshire, East Dunbartonshire, and North Lanarkshire) and Cardiff. Interesting, Greater Manchester, Merseyside, and Tyne and Wear are the areas with the highest enrichment of moderate local connectivity.

## Discussions

The main focus of the study is to provide a new data-driven descriptive analysis of personal genealogical history of a large modern population using IBD sharing patterns. By analyzing the 3.5 billion IBD segments >5 cM shared among half a million UK Biobank participants, our analysis offers a unique angle into the very recent demographic history of the United Kingdom. Unlike existing studies that focus on IBD segments between all individuals that reflect the history of the population ^3,8,9^, we are focusing on the personalized genealogical histories of individuals.

Based on approximately 14,000 IBD segments shared between a person and the rest of the UK Biobank participants, we discussed 3 categories of personalized genealogical descriptors: the genome coverage by IBD segments, the change of cousin counts at different IBD lengths, and the cousin enrichment within a neighborhood. Each of these genealogical descriptors reveals interesting details about personal genealogical history.

First, our analysis of genome coverage by IBD segments provides much higher resolution detail about the sharing of genetic information in a large modern population. We found even sampling the UK population at a rate of merely 0.7%, more than half of UK individuals have 80% of their genome reconstructable using IBD segments shared with others. Our analysis offered intricate implications as to genome privacy. As the fast growing business of direct-to-consumer (DTC) genomics, genome privacy has become of importance in both academia and the general public. Previously the concerns have been focused on the potential discrimination by employers and health insurance companies against carriers of certain mutations ^13,14^. Recently, a new concern ^15^ has been raised regarding law enforcement’s access to biobank scale genomic databases for solving criminal cases by connecting remote relatives genetically. Our study demonstrated that those two concerns are indeed two faces of the same coin as one’s genome can be largely reconstructed by the genomes of his/her genetic cousins traditionally thought as “unrelated”. Even sharing a low resolution of one’s genome, such as genotypes of 500K non-clinically-associated SNVs used by a DTC genomics company, may grant access to a much more detailed genomic information at base-pair resolution. Therefore, public awareness of this issue of genome privacy is needed.

Second, our analysis of the variation of cousin counts at different IBD lengths revealed individualized family expansion history. While some existing studies of population IBD patterns focused on clustering individuals based on the IBD network, we offered an individual-centered perspective. A unique benefit of our analysis is that instead of crawling the entire IBD network, such information can be collected efficiently via IBD queries such as PBWT-Query ^12^. For example, we identified a minority group of individuals (likely British Jews) that went through drastic recent expansion. Of interest for potential future research is to derive theoretical frameworks for inferring more details of an individual’s family history.

Third, we also demonstrated that IBD information of individuals can be intersected with other information, such as geographic location, and offer new measurement of social behaviors. Using cousins as a proxy of one’s extended family, we showed that by studying the enrichment of cousins in one’s neighborhood, an individual’s preference towards staying with extended family (local connectivity) is revealed. Interestingly, drastic regional variations of preferences of local connectivity were shown. In Greater London, people with weakest local connectivity are found, reflecting its cosmopolitan status. In Greater Manchester, people with moderate enrichment of local connectivity are found, indicating regional demographic movement in central England. In major cities of Scotland and Wales, people with strong local connectivity are found. By revealing regional genetic connections, our approach may open new research avenues of studying family-related social behaviors.

Our study is a population-scale population genetics study, i.e., the study sample size approaches the order of magnitude of the entire population. Traditional genetics studies are mostly at the sub-population scale, i.e., the samples under study were either a collection of “unrelated individuals” or familial members with traceable pedigrees. Population-scale data sets such as the UK Biobank offer opportunity of studying emergent phenomena that are not possible at the sub-population scale.

This analysis offers an alternative definition of ancestry. Traditionally, a person is typically labeled by one or a few predefined continental-level grouping. Rather than resorting to these somewhat arbitrary labels, we offer the geographic label of one’s ancestors, e.g., their birth locations, an idea gaining traction in recent literature ^16^. It is a well-known fact that the number of ancestors of each individual grows quasi-exponentially with the number of generations in the past. These ancestors may not have the same geographic label. Moreover, the distribution of ancestors’ geographic locations varies from generation to generation. Analyses of these patterns and theoretical modeling may be another exciting new research avenue.

## Methods

### IBD segment calling using RaPID

RaPID v.1.7 was run for all 22 autosomal chromosomes of all phased haplotypes in the UKBB comprising 487,409 participants and 658,720 sites ^1^. The parameters for RaPID were calculated assuming a genotyping error rate of 0.25%. The number of runs was set to 10, the minimum number of passes to 2, and the window sizes to 3. The minimum target length was set to 5 cM (-r 10 -s 2 -w 3 -l 5). The genetic maps from deCODE ^17^ for hg38 were downloaded and lifted over to h19 using the liftOver tool^18^. The longest monotonically increasing subset of the sites in each chromosome of hg19 was selected and subsequently interpolated to obtain the genetic locations of the available sites in the UKBB.

### Runs of PBWTs over genetic distance (RaPID v.1.7)

We modified RaPID to allow direct use of genetic distances in PBWT ^19^. The latest version of RaPID can take the minimum target length directly in cM and will return all detected segments greater than or equal to the given length without post-processing of data. The program holds a genetic mapping table for all the available sites and the PBWT has been modified to work directly with the genetic length instead of the number of sites. These changes were implemented in RaPID v.1.7.

We compared the results of the previous version of RaPID (v.1.2.3) and the new version (v.1.7) using the UK Biobank data. We found that the runtime has decreased from 12.78 days to 5.25 days, primarily due to the accurate control of window sizes.

### Benchmarking RaPID results versus KING

The relatedness data (up to third-degree relationship) from genotypes generated by KING were downloaded from the UK Biobank project ^1^. In order to distinguish parent/offspring and full siblings pairs in the first-degree relationships, an IBS0 cutoff of 0.002 was selected. If the IBS0 value was greater than 0.002, then the pair was considered as full siblings.

### Benchmarking RaPID results versus GERMLINE, Refined IBD and iLash

While it is possible to benchmark IBD segment detection methods using simulated data, it is almost impossible to capture all of the nuances in the real data. On the other hand, it is difficult to compare different methods within real data as there is often a lack of ground truth. We investigated the consistency among the IBD calls from different methods and then used the small subsets of MZ twin pairs for evaluation of detection power. The parameters for running GERMLINE and Refined IBD can be found in Supplementary Table S5.

We collected IBD results from four tools (GERMLINE, iLASH, RefinedIBD and RaPID) as four distinct sets of IBDs: S_1_, S_2_, S_3_, S_4_ respectively. All IBD segments that have been reported by at least one of the tools (in any of the result sets) were further investigated. To check whether an IBD1 in *S*_1_ is reported in *S*_2_, we searched in *S*_2_ for the same pair of individuals and checked if there is any reported IBD segment that overlaps at least 50% or more of the reported segment in *S*_1_. The percentage of IBD segments in each set that have been covered by other tools have been reported in the Supplementary Table S1.

### Estimation of detection power and accuracy

To estimate the detection power of different methods, 179 pairs of MZ twins (reported by KING) were considered. We expected that the reported IBD segments would cover the entire chromosomes between any two MZ twins. The detection power for each MZ twin pair was defined as the percentage of the genome (all 22 chromosomes) that has been covered by reported IBD segments. The average detection power values between all MZ twins among British, Non-British and all pairs were reported. The field 21000 from the UK Biobank data was used to determine the ethnic backgrounds of the MZ twins.

Overall, there are low chances that a false positive segment over 5 cM is called by any of these methods. In order to investigate the accuracy of the reported segment implicitly, 6225 parent/offspring pairs were used. Assuming there is no inbreeding, reported IBD segments between parent and offspring should not overlap. Please note that multiple IBD segments in each chromosome may be reported between an offspring and its parent due to recombinations or phasing error. We collected all 6276 parents/offspring pairs identified by KING. 51 pairs with total IBD length >3400 cM (reported by RaPID) were filtered out, which may be due to inbreeding. 382 pairs with total IBD length < 3000 cM (reported by RaPID) were also discarded, which might be due to low-quality phasing/genotyping. We searched for potential false positives, i.e., the overlapping IBD segments between parent/offspring pairs. An IBD segment is considered to be false positive if 50% of the segment is covered by another IBD segment between the same pair of parent/offspring. We find <0.01% potential false positives in any of these methods. If we defined the accuracy as percentage correctly identified segments: 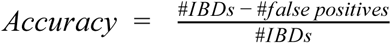, then the accuracy values for RaPID, GERMLINE, iLash and Refined IBD using parent/offspring pairs would be 99.9032%, 99.9286%, 99.95813% and 99.9512%, respectively. Manual inspection of false positives also revealed that most cases are also likely due to runs-of-homozygous segments.

### Assignment of geographic area

Ordinance coordinates in the form of (east, north) were retrieved from the UKBB fields 22702 and 22704. The coordinates were converted to longitudes and latitudes using the Python library *convertbng* (https://pypi.org/project/convertbng/). The Python library *reverse_geocoder* (https://pypi.org/project/reverse_geocoder/) was then used to find the nearest town/city using the GPS coordinates. For each coordinate, *reverse_geocoder* returns only the corresponding subdivision, which is finer than counties. The subdivisions were then translated into the corresponding counties.

### Cousin count adjustment by the regional sampling rate

The population size for each county was extracted ^20^. For each county *i*, the sampling rate *S*_*i*_ was defined as the number of participants divided by the population size. For each individual, the number of cousins in each county (*C*_*i*_) was calculated (using the home locations). The normalized number of cousins for each individual was then calculated using the following formula: 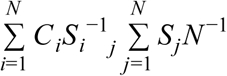, where *N* denotes the total number of counties. 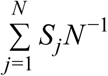 simply denotes the average sampling rate for all counties.

### Searching for query haplotype from the Personal Genome Project in UKBB

The trio data of an Ashkenazi Jewish family (huAA53E0 son and his parents hu8E87A9, hu6E4515) were downloaded ^21^. The sites containing SNPs were extracted from the trio data. Each site containing a missing value was discarded. The genotype data for the son were then phased using parent data and simplified by filtering out all-het sites. The overlapping sites with the UKBB (chromosome 1) were selected which resulted in 6629 sites. The 6629 sites of chromosome 1 for all the individuals from UKBB were extracted using vcftools (v0.1.15) ^22^. PBWT-Query ^12^ was used to search for the query haplotypes with a minimum target length of 200 SNPs (-L 200).

### Design of genetic genealogical descriptors

The following categories of genetic genealogical descriptors were defined in order to capture extensive information from the cousin cohorts of an individual: 1) Percentage of the genome covered by IBD segments from cousins, 2) Decay of cousin counts as the length of IBD segments increase, 3) Enrichment of cousins in one’s neighborhood.

### Enrichment of counts in tables

For describing the enrichment of counts of individuals in a contingency table, e.g., e25 vs city_size or c vs area, we used the Pearson’s residual: ((observed − expected)/sqrt (expected)).

### Percentage of the genome covered by IBD segments

In order to investigate the correlation between the available number of individuals from a population and the percentage of the genome covered, 10 subsets of the UK Biobank were extracted. The largest available population contained 480,518 individuals. 9 other subsets were extracted by randomly selecting half of the individuals from each subset, subsequently. For example, the next subset contained 250,814 individuals. For each subset, the average genome coverage was computed as follows: Each chromosome was divided into bins of 1 Mbps. For each individual, the number of shared IBD segments overlapping with each bin was calculated. Then, the number of bins overlapping with at least 1, 5, 10 IBD segments were calculated and divided by the total number of bins. Finally, the average of the genome coverage among all available individuals was computed.

### Collecting IBD segment counts at different lengths

For each individual, counts of cousins sharing IBD segments at different lengths are informative to personal genealogical history. We chose the counts of IBD segments in [5, 10), and [10, 3400). The total sum of IBD segments between any two pairs of individuals sharing an IBD segment (cousins) from all autosomal chromosomes in the UKBB was calculated. Subsequently, the number of cousins of each individual for two bins (5-10 cM and >= 10 cM) was computed.

### Enrichment of cousins in one’s neighborhood

The UK Biobank home location fields 22702_0_0 (east coordinate) and 22704_0_0 (north coordinate) were used to compute the distance between any two individuals. The number of neighbors and cousins within a neighborhood of 1, 5, 10 and 25 km were calculated as follows: For each individual and the given radius *r* (e.g. 1km), all UK participants within a distance *r* from both query’s east and north coordinates were extracted. Subsequently, the Euclidean distances between the query individual and the extracted individuals were calculated.

### Designating types of living environment by counting neighbors

By plotting the histogram of the number of neighbors within 25km (Figure 4b), we divided the individuals as dwellers of “Metropolitan”, “Big City”, “Mid-sized City”, “Small City”, and “Rural”, as people with count of neighbors included in UK Biobank >40000, 31000-40000, 21000-31000, 5000-21000, and <5000, respectively.

## Acknowledgments

A.N., S.Z., and D.Z. were supported by the National Institute of Health (grant no. R01 HG010086). X.L. was supported by the National Institute of Health (grant no. grant R01 HG009524). This research has been conducted using the UK Biobank Resource under Application Number 24247.

---

**Supplementary Figure S1:**
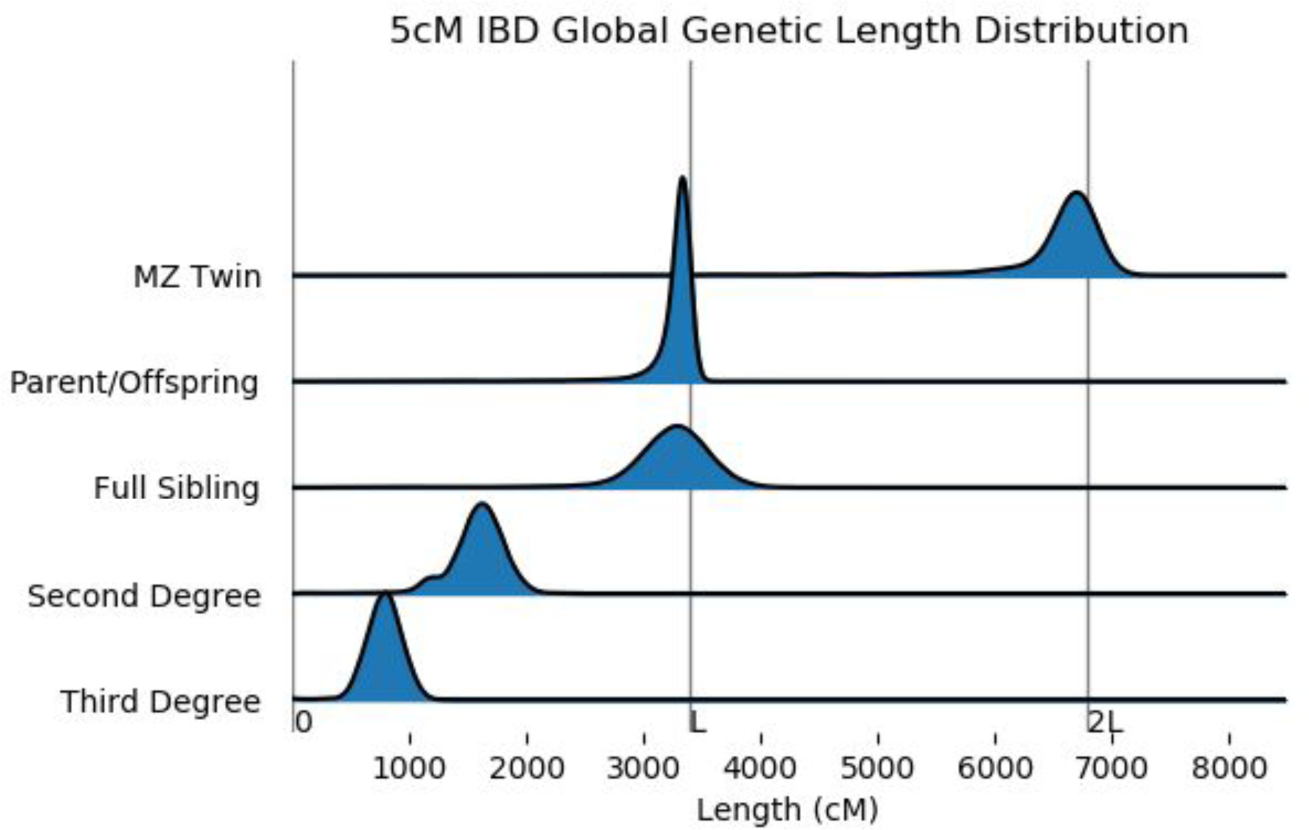
Distribution of kinship coefficients derived from RaPID’s result is consistent with expected value in close relative pairs identified by KING.

**Supplementary Figure S2:**
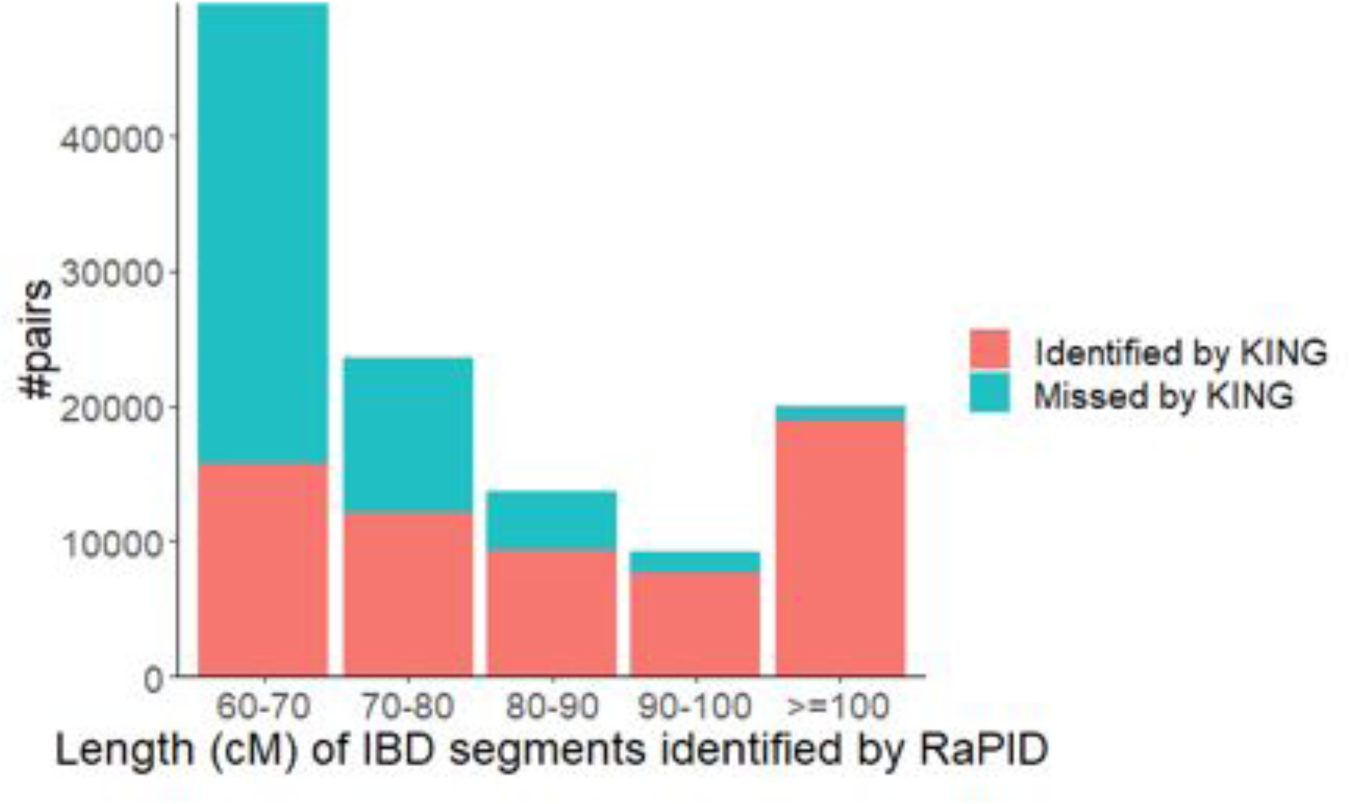
A large fraction of IBD segments identified by RaPID are not among the 3rd degree related pairs.

**Supplementary Figure S3:**
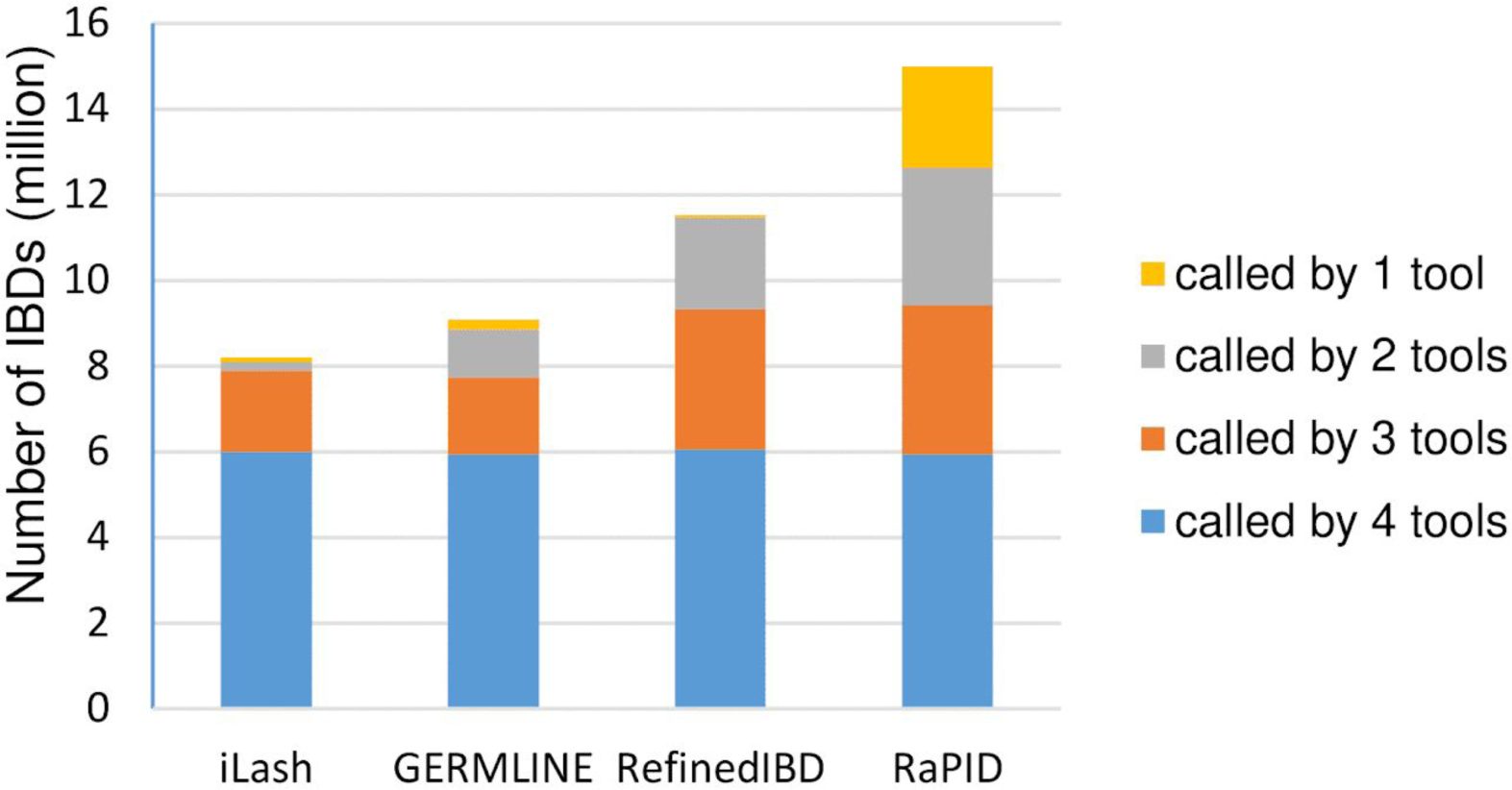
Number of IBD calls for different IBD detection tools using chromosome 22 of 200K individuals from UK Biobank. IBD calls for each tool are stratified by the total number of tools that identified the segments.

**Supplementary Figure S4:**
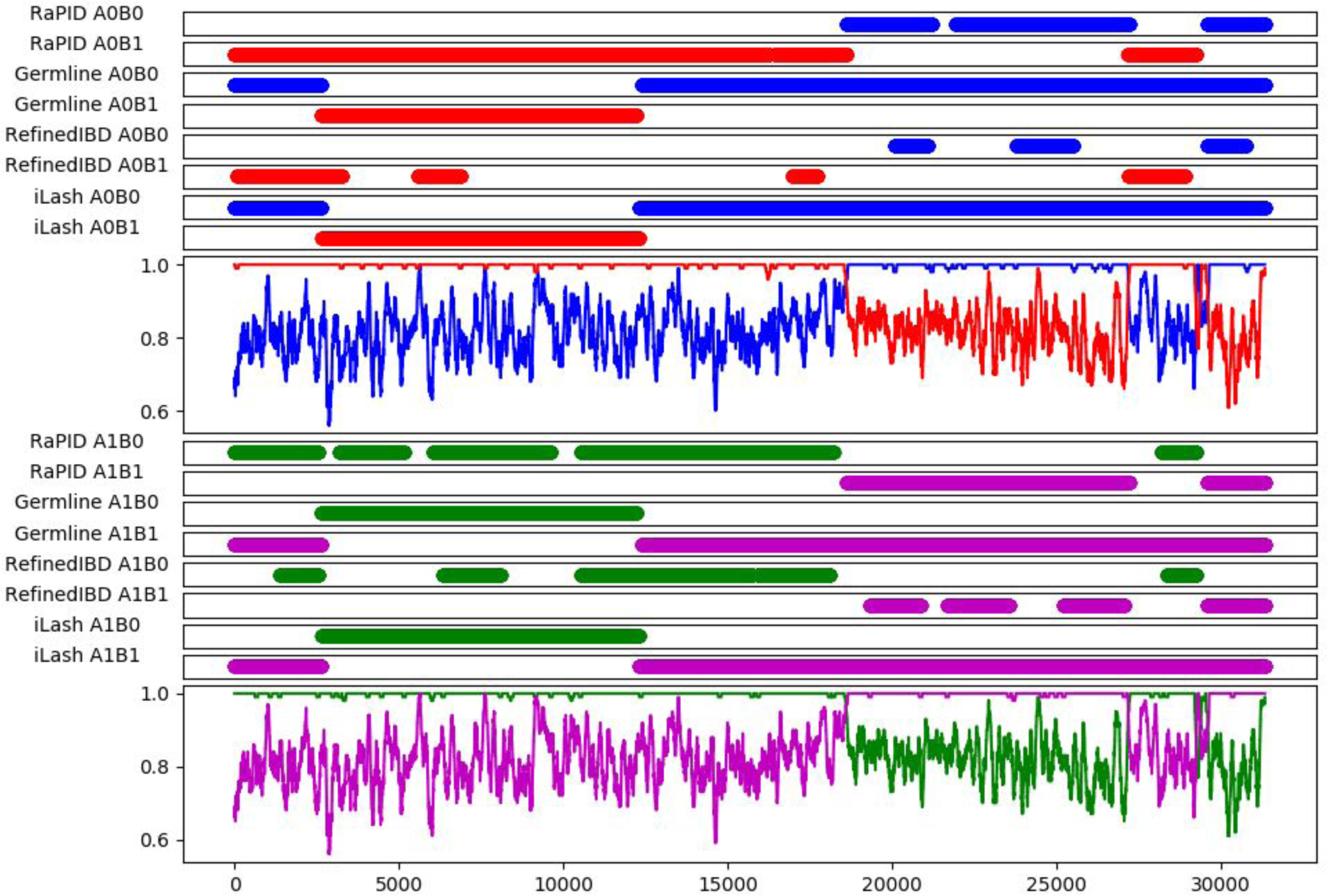
An example of IBD segments over chromosome 12 called by different methods using a twin pair. The identities of matches over the moving averages of windows with 100 SNPs were computed using four different combinations of haplotypes: A0B0, A0B1, A1B0, and A1B1. A0B0 matches are represented in blue, A0B1 in red, A1B0 in green, and A1B1 in purple. IBD results for each tool have been depicted on the top tracks. The x-axis denotes the site index of chromosome 12.

**Supplementary Figure S5:**
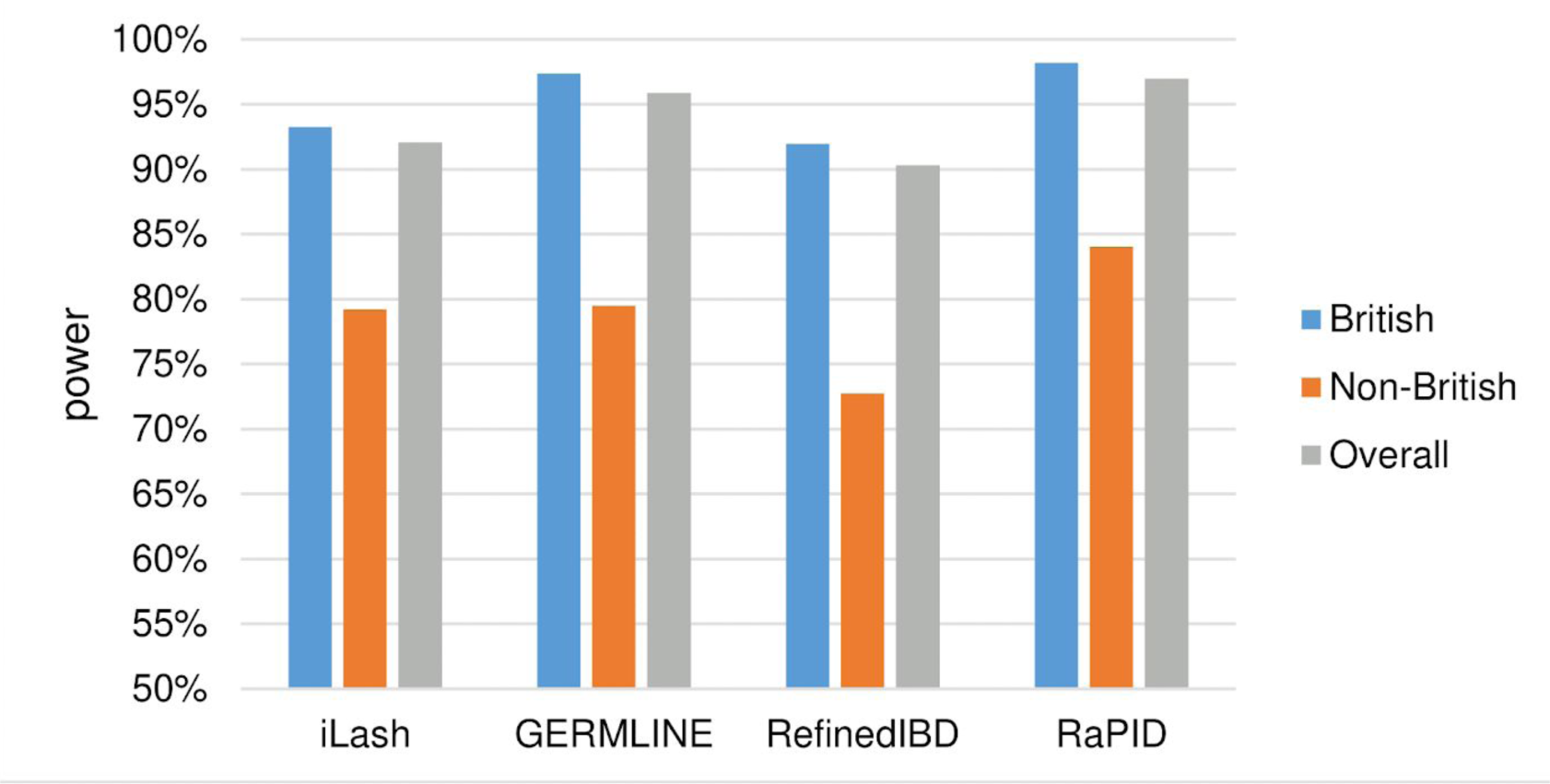
Average detection power of different methods.

**Supplementary Figure S6:**
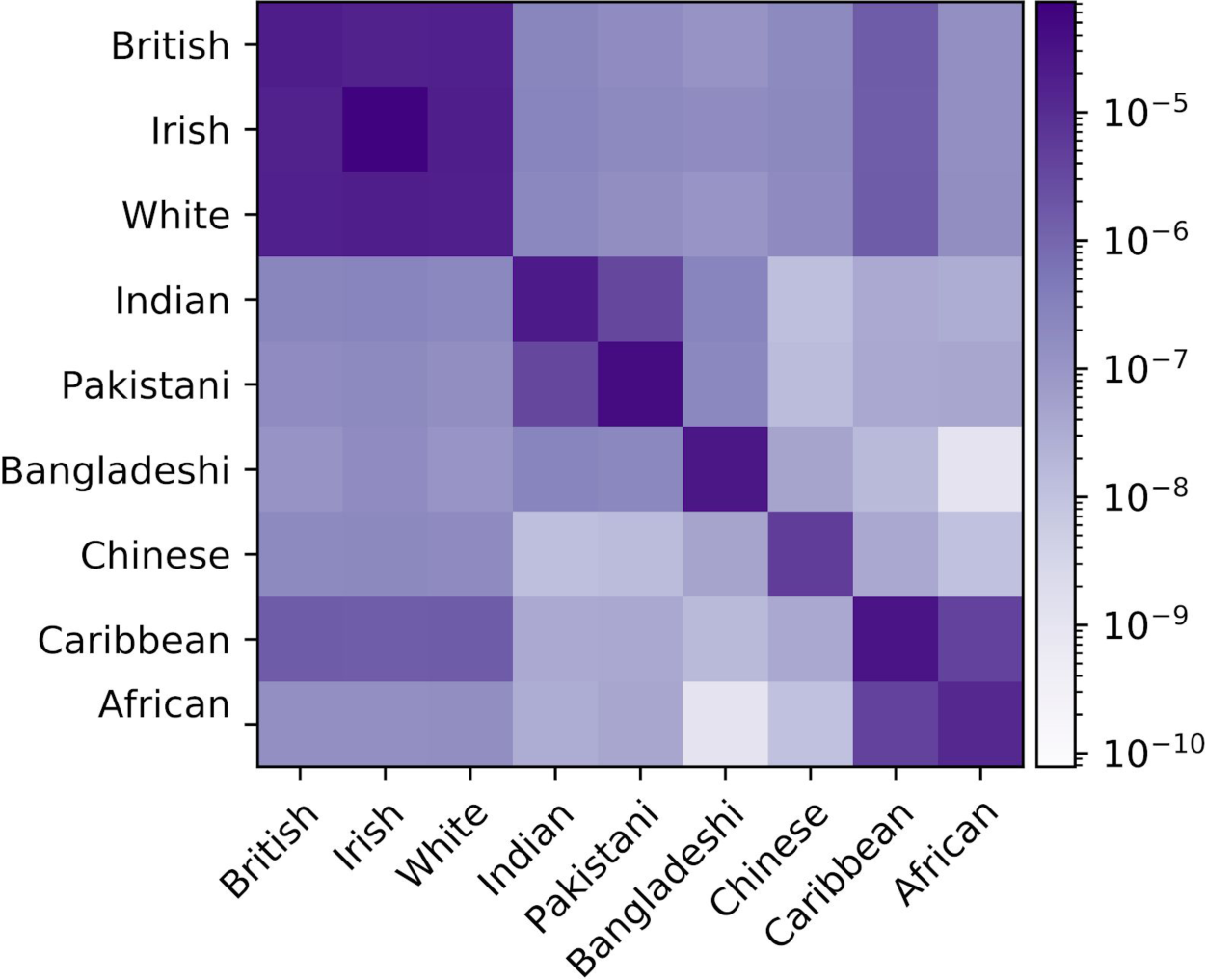
Ethnicity by ethnicity kinship matrix using the self-reported ethnic background in UK Biobank. Intrapopulation kinship values are higher than interpolation kinship values, as expected. The kinship value between British and Irish is high, however the Irish population has a distinguishable intrapopulation kinship. The closest population to Indian is Pakistani according to the kinship values. The African population has the highest kinship with the Caribbean among the other ethnic groups as anticipated.

**Supplementary Figure S7:**
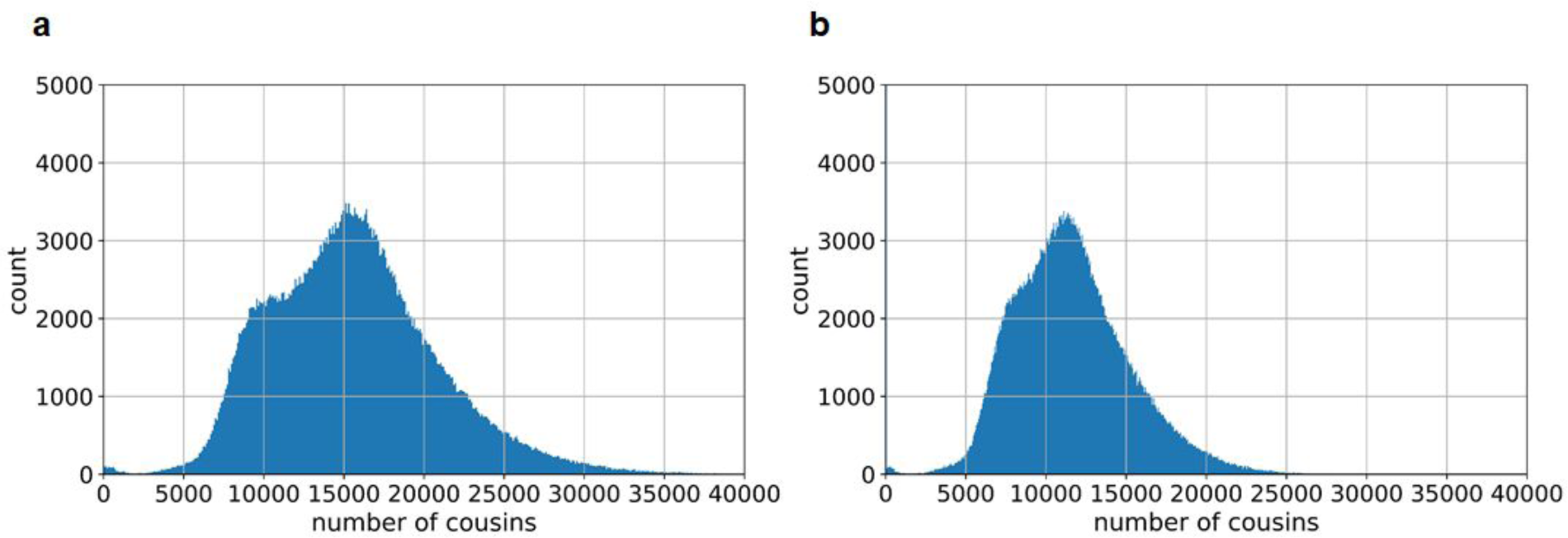
Cousin count of British. (a) unadjusted; (b) adjusted by regional sampling rates.

**Supplementary Figure S8:**
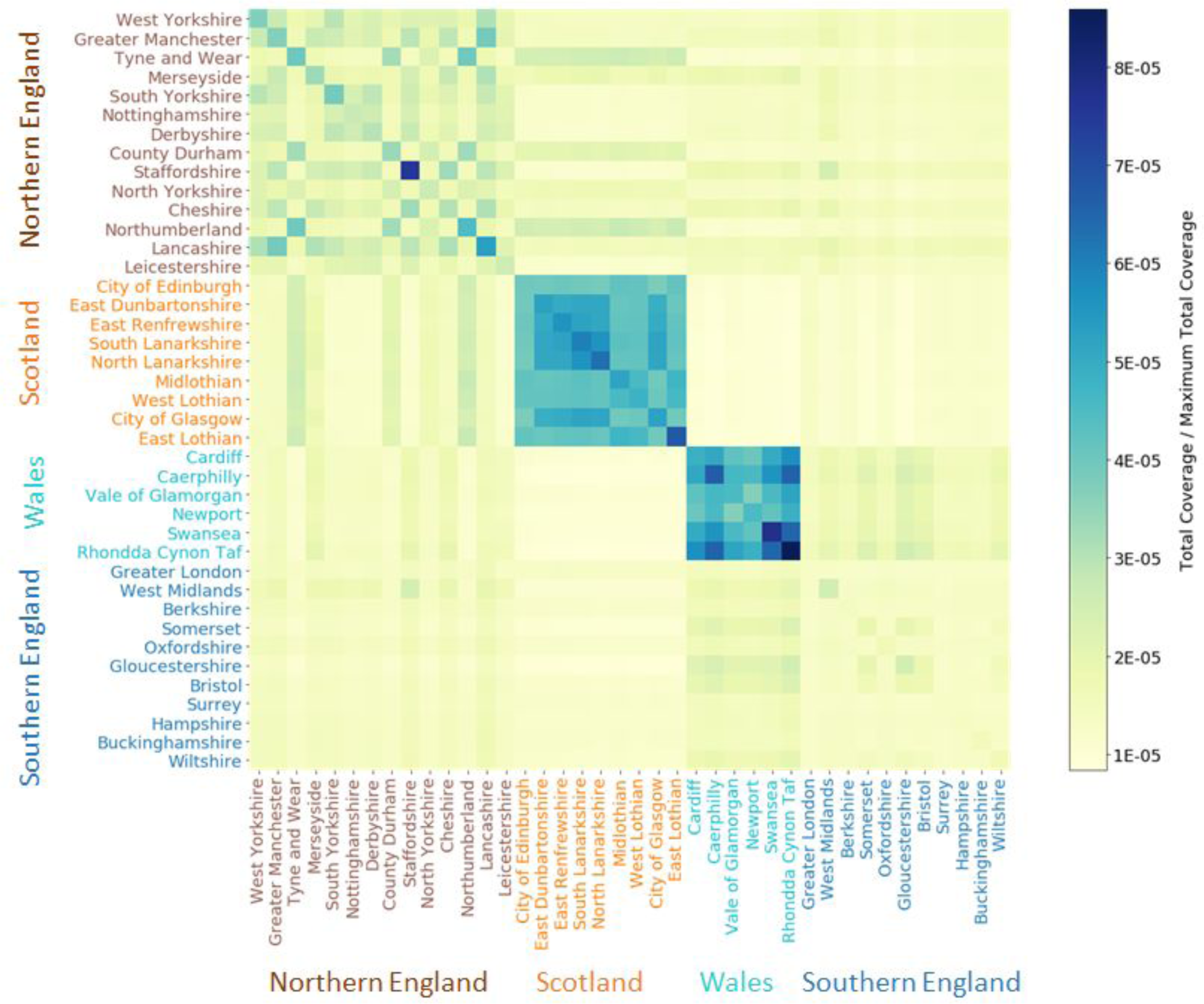
Cross-region average cousin count. Numbers are normalized by total potential pairs and the total length of the chromosomes.

**Supplementary Figure S9:**
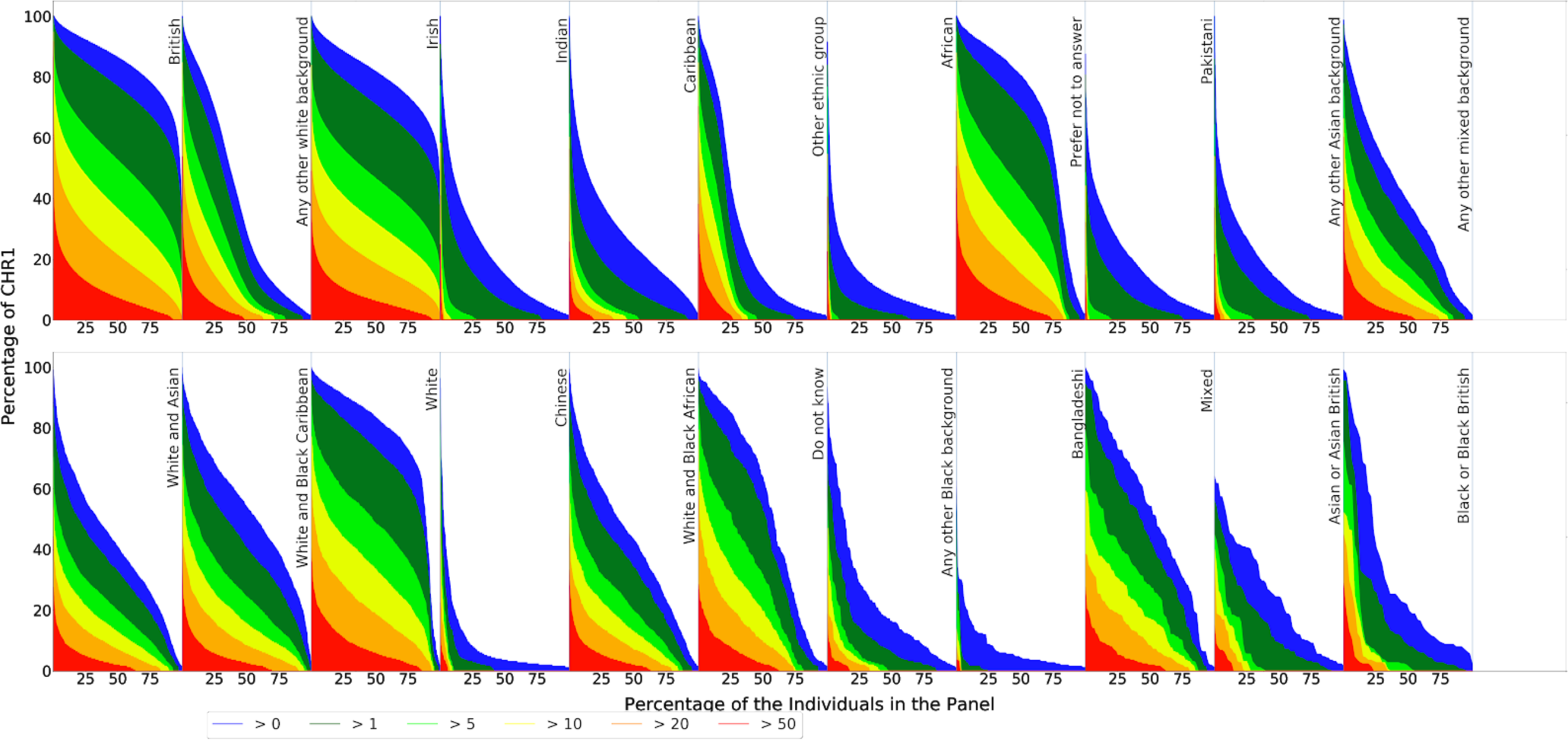
Percentage of the genome covered by IBD segments from others in UK Biobank by ethnicity.

**Supplementary Figure 10:**
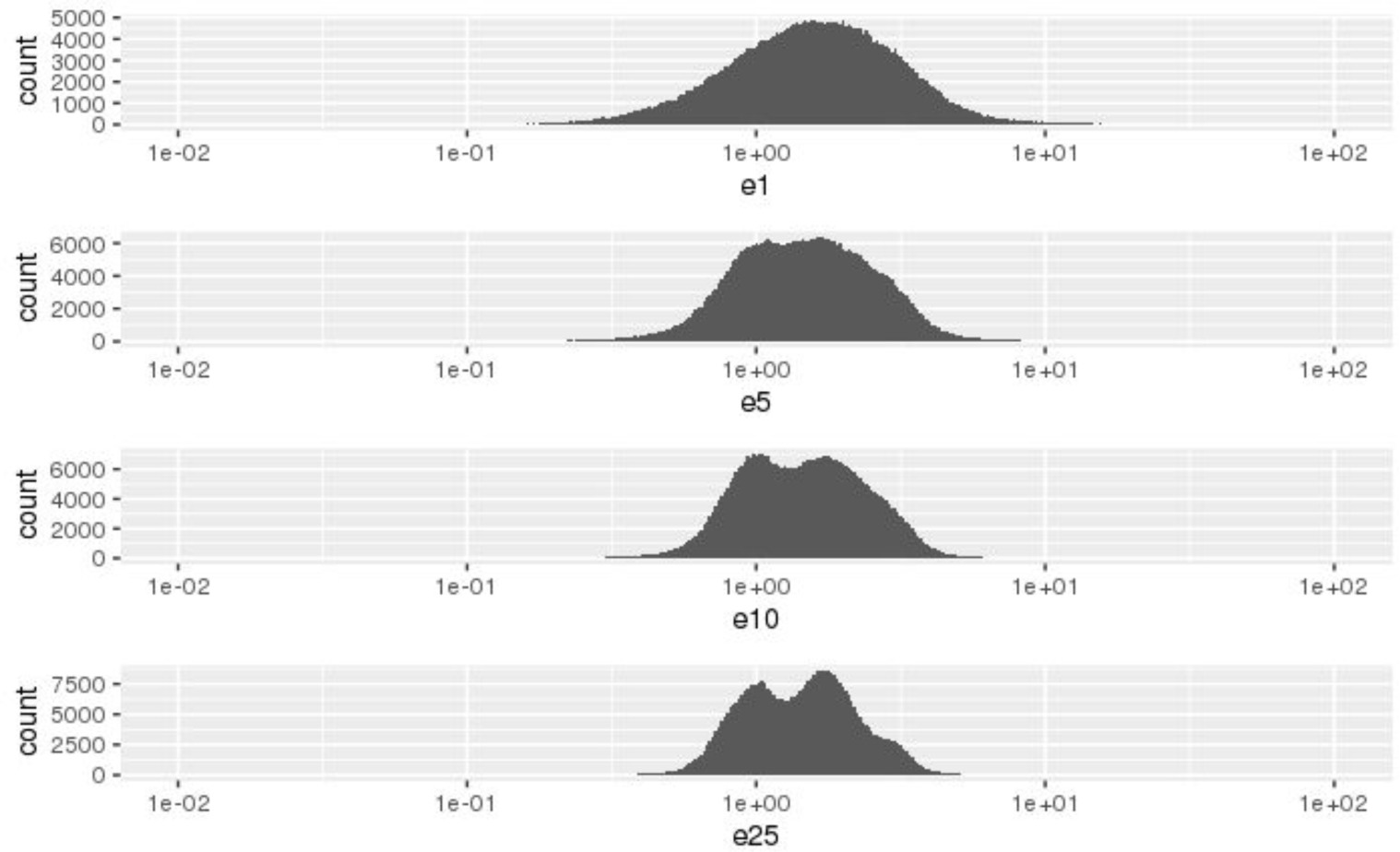
The distribution of CEIN of all UK Biobank participants.

**Supplementary Table S1:** IBD results of Germline, iLash, RefinedIBD and RaPID (rows) covered by other tools (columns).

|  | Covered by<br>GERMLINE | Covered by<br>iLASH | Covered by<br>RefinedIBD | Covered by<br>RaPID |
| --- | --- | --- | --- | --- |
| **GERMLINE** |  | 68.53% | 82.51% | 95.76% |
| **iLASH** | 77.27% |  | 93.61% | 97.24% |
| **RefinedIBD** | 66.60% | 67.13% |  | 99.45% |
| **RaPID** | 58.49% | 52.75% | 75.57% |  |

**Supplementary Table S2:** Average detection power of different methods.

|  | British<br>(n=164) | Non-British<br>(n=15) | Overall<br>(n=179) |
| --- | --- | --- | --- |
| **RaPID** | 98.17% | 84.03% | 96.98% |
| **GERMLINE** | 97.36% | 79.48% | 95.86% |
| **RefinedIBD** | 86.02% | 63.97% | 84.18% |
| **iLash** | 93.23% | 79.20% | 92.06% |

**Supplementary Table S3:** Cousin enrichment in different neighborhoods decays with increasing radius.

| radius (km) | 1 | 5 | 10 | 25 |
| --- | --- | --- | --- | --- |
| average #neighbors | 295 | 2851 | 8438 | 26528 |
| average cousin enrichment | 1.703 | 1.650 | 1.613 | 1.588 |

**Supplementary Table S4:** Enrichment of people with different local connectivities in UK areas, as indicated by Pearson’s residuals.

| Area | Weak | Moderate | Strong |
| --- | --- | --- | --- |
| Aberdeen City | -0.1 | -3.4 | 8.8 |
| Aberdeenshire | 1.6 | -2.5 | 2.5 |
| Angus | 0.8 | -2 | 3.3 |
| Argyll & Bute | 1.1 | -3.3 | 5.8 |
| Avon | 27.5 | -16.1 | -22.8 |
| Bedfordshire | 5.8 | -4.5 | -1.9 |
| Berkshire | 92.7 | -70.3 | -36.5 |
| Blaenau Gwent | -1.7 | -0.8 | 6 |
| Bridgend | -1.2 | -1.2 | 5.7 |
| Buckinghamshire | 30.5 | -23.3 | -11.5 |
| Caerphilly | -23.1 | -15.5 | 92.1 |
| Cambridgeshire | 6.2 | -4.9 | -1.9 |
| Cardiff | -20.4 | -12.5 | 78.6 |
| Carmarthenshire | -1.9 | -3.4 | 12.8 |
| Ceredigion | 2.6 | -3 | 1.5 |
| Cheshire | -13.8 | 19.1 | -16.2 |
| Clackmannanshire | -1.1 | -1.4 | 5.9 |
| Conwy | 1.8 | -2.2 | 1.4 |
| Cornwall and Isles of Scilly | 5 | -5.7 | 2.8 |
| Cumbria | 4.6 | -4.5 | 0.9 |
| Denbighshire | 0.6 | -0.9 | 1 |
| Derbyshire | -9.4 | 23 | -36.2 |
| Devon | 7.2 | -6.9 | 0.7 |
| Dorset | 4.3 | -4.4 | 1.1 |
| Dumfries & Galloway | 1.5 | -3.1 | 4.4 |
| Dundee City | 0.5 | -3.1 | 6.6 |
| Durham | -25.6 | 33.2 | -24.6 |
| East Ayrshire | -2.1 | -0.6 | 6.4 |
| East Dunbartonshire | -30.5 | -21.7 | 124.6 |
| East Lothian | -15.2 | 0.3 | 34.2 |
| East Renfrewshire | -30.3 | -21.3 | 123.3 |
| East Sussex | 6.2 | -5.1 | -1.6 |
| Edinburgh, City of | -34.9 | -4.3 | 91.1 |
| Eilean Siar | 1.4 | -2.2 | 2.5 |
| Essex | 7.8 | -6.1 | -2.7 |
| Falkirk | -3.1 | -2 | 12.3 |
| Fife | -4.3 | -0.5 | 11 |
| Flintshire | -3.1 | 3.8 | -2.3 |
| Glasgow City | -21 | -14.9 | 85.7 |
| Gloucestershire | 4.1 | 7.8 | -29.2 |
| Greater London | 163 | -127.8 | -54.4 |
| Greater Manchester | -57.3 | 72.2 | -49.9 |
| Gwynedd | 2.2 | -3.3 | 3.2 |
| Hampshire | 49.6 | -37.6 | -19.7 |
| Hereford and Worcester | 1.2 | 2 | -7.9 |
| Hertfordshire | 12.2 | -9.4 | -4.5 |
| Highland | 2.6 | -4 | 4.1 |
| Humberside | 5.7 | -5 | -0.7 |
| Inverclyde | -2.5 | -1.5 | 9.5 |
| Isle of Anglesey | 2.4 | -2.2 | 0 |
| Isle Of Wight | 3.2 | -2.5 | -1 |
| Kent | 9.8 | -8.1 | -2.3 |
| Lancashire | -28.1 | 26.5 | -2.1 |
| Leicestershire | 13.4 | -6.8 | -13.6 |
| Lincolnshire | 6.9 | -5.8 | -1.2 |
| Merseyside | -49.6 | 61.1 | -39.6 |
| Merthyr Tydfil | -3.8 | -1.2 | 11.7 |
| Midlothian | -22.5 | 0.6 | 50.2 |
| Monmouthshire | 4.7 | -2.3 | -5.1 |
| Moray | 1.1 | -2.5 | 3.8 |
| Neath Port Talbot | -6.5 | -8.9 | 37.5 |
| Newport | -12.2 | 0.5 | 26.8 |
| Norfolk | 5.2 | -5.2 | 1.3 |
| North Ayrshire | -1.6 | -2.5 | 9.9 |
| North Lanarkshire | -30.2 | -20.5 | 120.9 |
| North Yorkshire | 9.8 | 2.6 | -29.2 |
| Northamptonshire | 6.4 | -4.9 | -2.4 |
| Northumberland | -23.8 | 19.6 | 5.3 |
| Nottinghamshire | 6.9 | 8.8 | -38 |
| Orkney Islands | 1.6 | -1.2 | -0.5 |
| Oxfordshire | 75.5 | -56.2 | -32.4 |
| Pembrokeshire | 3.2 | -3.3 | 0.7 |
| Perth & Kinross | 1.1 | -2.8 | 4.4 |
| Powys | 2.4 | -2.6 | 1 |
| Renfrewshire | -15.1 | -9.9 | 59.7 |
| Rhondda, Cynon, Taff | -16.4 | -12.7 | 69.7 |
| Rutland UA | 2.3 | -1.6 | -1.3 |
| Scottish Borders | 1.4 | -1.9 | 1.7 |
| Shetland Islands | 1.3 | -1 | -0.4 |
| Shropshire | 2.3 | -3.3 | 3 |
| Somerset | 24.1 | -8.5 | -34.2 |
| South Ayrshire | -0.4 | -1.3 | 4.1 |
| South Lanarkshire | -32.9 | -21.7 | 130.3 |
| South Yorkshire | -38.6 | 50.4 | -38 |
| Staffordshire | -45.6 | -12.5 | 136.3 |
| Stirling | -4 | -2.7 | 16 |
| Suffolk | 4.4 | -3.8 | -0.5 |
| Surrey | 65.2 | -50.8 | -22.2 |
| Swansea | -11.7 | -19.1 | 74.9 |
| The Vale of Glamorgan | -8.9 | -8.9 | 42.8 |
| Torfaen | -2.2 | -1.8 | 9.7 |
| Tyne And Wear | -60.2 | 60.5 | -13.8 |
| Warwickshire | 2.5 | 0.7 | -7.4 |
| West Dunbartonshire | -12.9 | -7.5 | 48.6 |
| West Lothian | -22.4 | 2.3 | 45.8 |
| West Midlands | 5.5 | 9.4 | -36.3 |
| West Sussex | 6.6 | -5.5 | -1.4 |
| West Yorkshire | -38.4 | 56.2 | -52.9 |
| Wiltshire | 15.3 | -10.4 | -8.9 |
| Wrexham | -7.3 | -6.9 | 34 |

**Supplementary Table S5:** Parameters and command lines for benchmarking different IBD detection tools.

|  | Parameters | Command Line |
| --- | --- | --- |
| GERMLINE | -min_m 5<br>-h_extend<br>-haploid | ./germline -input ukb_22_200k.vcf.ped<br>ukb_22_200k.vcf.map -output<br>ukb_22_200k_5cm -min_m 5 -h_extend<br>-haploid |
| RefinedIBD | length=5 | java -Xmx200g -jar<br>refined-ibd.12Jul18.a0b.jar<br>gt=ukb_22_200k.vcf out=22_200k<br>map=decLift_HG19_22.map length=5 |
| iLash | slice_size 350<br>step_size 350<br>perm_count 20<br>shingle_size 15<br>shingle_overlap 0<br>bucket_count 5<br>max_thread 20<br>match_threshold<br>0.99<br>interest_threshold<br>0.70<br>max_error 0<br>min_length 5<br>auto_slice 1<br>cm_overlap 1<br>minhash_threshold<br>55 | ./IBD iLa_22.config |
| RaPID | -w 3 -r 10 -s 2 -d<br>5 | ./RaPID_v.1.7 -i 22.vcf.gz -o 22_5 -w 3<br>-r 10 -s 2 -d 5 -g 22.rMap |

